# Complete list of canonical post-transcriptional modifications in the *Bacillus subtilis* ribosome and their link to RbgA driven large subunit assembly

**DOI:** 10.1101/2024.05.10.593627

**Authors:** Anna M. Popova, Nikhil Jain, Xiyu Dong, Farshad Abdollah-Nia, Robert A. Britton, James R. Williamson

## Abstract

Ribosomal RNA modifications in prokaryotes have been sporadically studied, but there is a lack of a comprehensive picture of modification sites across bacterial phylogeny. *B. subtilis* is a preeminent model organism for gram-positive bacteria, with a well-annotated and editable genome, convenient for fundamental studies and industrial use. Yet remarkably, there has been no complete characterization of its rRNA modification inventory. By expanding modern MS tools for the discovery of RNA modifications, we found a total of 25 modification sites in 16S and 23S rRNA of *B. subtilis,* including the chemical identity of the modified nucleosides and their precise sequence location. Furthermore, by perturbing large subunit biogenesis using depletion of an essential factor RbgA and measuring the completion of 23S modifications in the accumulated intermediate, we provide a first look at the order of modification steps during the late stages of assembly in *B. subtilis*. While our work expands the knowledge of bacterial rRNA modification patterns, adding *B. subtilis* to the list of fully annotated species after *E. coli* and *T. thermophilus,* in a broader context, it provides the experimental framework for discovery and functional profiling of rRNA modifications to ultimately elucidate their role in ribosome biogenesis and translation.

## INTRODUCTION

RNA components of the ribosome, an essential ribonucleoprotein machine, are extensively decorated with post-transcriptional rRNA modifications, among which methylations of the base or ribose, and pseudouridines are the most frequent. rRNA is modified during the ribosome biogenesis process using either site-specific protein enzymes or an RNA-guided mechanism present in archaea and eukaryotes. There is little conservation of modified sites among the three domains of life (1), with an increased number of modifications found in eukaryotic ribosomes (36 sites in *E. coli* vs 228 in mammalian cells (2)). The abundance of modification events adds to the overall complexity of the ribosome assembly process, and both modification machinery and modified nucleosides arguably play active roles in rRNA folding and protein binding steps during the assembly (3,4). Furthermore, many modifications are located in functionally important sites of the ribosome including the decoding center, peptidyl transferase center (PTC), peptide-exit tunnel, and inter-subunit bridges (5), where they presumably modulate local structure and interactions to ensure accurate and efficient protein synthesis.

The rRNA modification machinery remains an important target for the constantly evolving drug resistance in pathogenic organisms. Not only can an acquired modification be effective at preventing binding of antibiotics to their ribosome target sites, but the lack of a modification due to loss of function mutations can decrease susceptibility to drugs (6). While roles of individual modifications and corresponding enzymes in ribosome biology and in bacterial evolution are not well understood, the variability of rRNA modification patterns between species is evident and may provide further clues for functional investigations (SI Table 1).

Although rRNA is the most abundant RNA in cells, attempts to obtain a complete list of bacterial rRNA modifications have been surprisingly infrequent. *E. coli* and *T. thermophilus* are the only two organisms for which both 16S and 23S modification have been characterized in full. In *E.coli*, a set of 36 rRNA modifications has been reported and independently proven by many labs (3,5,7), and a total of 25 modified nucleosides placed in rRNA of *T. thermophilus* (8–10) (SI Table 1). Complete or nearly complete maps of 16S modifications have been reported for *T. maritima* (11), *C. acetobutylicum* (12), and *L. pneumophila* (13). Significant number of residues were discovered in rRNA of *C. sporogenes* (14), in *P. aeruginosa* (15), *S. aureus* (16), and in *B. subtilis*, where a total of ten rRNA methylations and pseudouridines were experimentally confirmed (17–22). In addition, many experimentally verified rRNA modification sites remain uncatalogued in RNA modification databases like Modomix (23) due to the fact that only a few were found per species (24–27).

In the era of Next-Generation (NGS) and Direct RNA Sequencing (DRS), Mass Spectrometry (MS) continues to play a crucial role in the discovery, independent validation, and quantitative profiling of modifications in abundant cellular RNAs like rRNA and tRNA. In fact, MS offers direct simultaneous identification of all the types of modified residues that produce a detectable mass shift even when placed in the vicinity of other modification sites. Unlike eukaryotic rRNA, where ribose methylations and pseudouridines are dominating the pool of modified nucleosides, bacterial rRNA is especially rich in diverse base modifications all of which may be difficult or impossible to detect using sequencing approaches alone. Furthermore, recent advances increasing MS instrumentation speed and sensitivity, improving bioinformatics analysis (28) and ribonuclease cleavage specificity (29,30) make MS-based discovery more efficient. Another advantage of MS is the opportunity for relative and absolute quantitation of modifications which is key for studying biological functions.

rRNA modifications are an integral part of the assembly process where binding and release of modification enzymes are tightly coordinated with structural rearrangements within the growing precursor. The transient nature of rRNA-modification enzyme interactions and low abundance of the assembly intermediates (∼ 2% of the total ribosome load in the bacteria cell) in general, make the modification steps difficult to study. Fortuitously, several cryo-EM structures have been reported with modification enzymes bound to their substrates, yet all the findings are currently limited to the late assembly stages (31–34). Perhaps, a more tangible approach is a direct observation of rRNA modifications in intermediate particles accumulated as a result of genetic perturbations, environmental stresses, and inhibition of assembly. Since single-particle EM resolution of 3-5 LJ is hardly enough to confidently assign modification status directly within the intermediate isolates, RNA MS offers a much more robust alternative to quantitative assessment, as previously demonstrated by our laboratory for *E. coli* rRNA modifications and assembly intermediates (35,36).

In this report, we expand our MS toolset to provide the complete set of modifications in 16S and 23S RNA of *B. subtilis*. Furthermore, we characterize the modification status of the abundant large subunit intermediate particles (45S^RbgA^) isolated from *B. subtilis* cells depleted of an essential ribosome biogenesis factor called RbgA. The 45S^RbgA^ intermediate has a scaffold of a nearly complete 50S subunit, but the A, P and E tRNA binding sites are unstructured (37). Quantitative inventory of RNA modifications using isotope ratio MS demonstrates that except for several positions in helix 69, all 23S modifications are fully incorporated into 45S^RbgA^. The missing modifications are installed by the homologs of pseudouridine synthase RluD and methyltransferases TlyA and RlmH, which are known to be late-acting protein enzymes in other bacteria species (34,36,38). These observations suggest that despite the highly flexible parallel nature of the assembly pathway, modification enzymes elicit their functions along the route strictly cooperating with the folding of the large structural blocks composed of rRNA and r-proteins. This precision is important to warrant complete modification status in matured 70S.

## MATERIALS AND METHODS

### *B. subtilis* growth, metabolic labeling, and rRNA purification

*B. subtilis* strain RB1202 was constructed by transforming chromosomal DNA from RB301 (39) to prototrophic py79 strain containing *rbgA* (a.k.a. *ylqf*) under the IPTG inducible P_spank_ promoter. For identification of modified nucleosides, RB1202 was grown in the MSpitz9 defined medium (40) using 1 mM IPTG for induction that leads to the synthesis of mature 50S and 70S ribosomal particles. To elicit specific changes to the mass of RNA and facilitate the discovery of RNA modifications, metabolic precursors containing stable isotopes were added. Particularly, 1 g/L of either (NH_4_)_2_SO_4_ (designated as ^14^N throughout the text) or (^15^NH_4_)_2_SO_4_ (designated as ^15^N, Cambridge Isotope Laboratories) was used as a source of nitrogen. C[^2^H]_3_-methionine (Cambridge Isotope Laboratories) labeling was carried out by adding deuterated methionine at a concentration of 50 mg/L directly to the ^14^N medium. For pseudouridines identification, 25 mg/L of 5,6-[^2^H]-uracil (CDN isotopes) precursor was added to both ^14^N and ^15^N labeled medium. All the bacteria cultures were incubated at 37°C until reached 0.5 OD_600_. Cells were harvested by centrifugation for 10 min at 5000 *g*.

To isolate ribosomes and separate 30S and 50S rRNA subunits, cells were lysed in the lysis buffer (10 mM Tris–HCl pH 7.5, 60 mM KCl, 10 mM MgCl_2_, complete EDTA-free protease inhibitor (Roche), 10 μg/ml RNase-free DNase (Qiagen), and 1 uL/ml of RNaseOUT (Thermo Fisher), 0.005% Tween20) using French press at 1400-1600 psi. Cell lysate was first clarified by centrifuging at 16 000 g for 20 minutes, then the ribosome pellet was isolated by centrifuging the lysate at 135 000 g for 2.25 h at 4°C. The pellet was gently washed using high salt buffer (20 mM Tris-HCl, pH 7.5, 10 mM MgCl_2,_ 800 mM NH_4_Cl, 1 mM DTT) and resuspended in 20 mM Tris-HCl, pH 7.5, 30 mM NH_4_Cl, 1 mM MgCl_2,_ 1 mM DTT. 18-43% (wt/vol) sucrose density gradient was made in 20 mM Tris-HCl, pH 7.5, 1 mM MgCl_2,_ 50 mM NH_4_Cl, 1 mM DTT. Ribosome resuspension was layered on the surface of the gradient and centrifuged for 14 hours at 21 000 rpm (∼ 88 000 g) and 4°C in the Surespin 630 rotor (Sorvall). Gradient fractions were collected using Biorad FPLC by monitoring OD_254._ Fractions assigned to the individual ribosomal subunits were pooled, then concentrated and buffer exchanged to 10 mM Tris-HCl, pH 7.5, 1 mM MgCl_2,_ 60 mM KCl,1 mm DTT using a 100 kDa Amicon filter device.

For quantitative analysis of modifications in RbgA depleted ribosomes, RB1202 cells were grown in MSpitz9 using 5 or 10 µM IPTG induction. For comparison, the doubling time at 5 µM IPTG induction was 111.7 min, at 10 µM IPTG - 78.4 min, and at 1 mM IPTG - 52.3 min. Cells were harvested and 45S^RbgA^ and 70S^RbgA^ particles were isolated via 10-40% sucrose gradient ultracentrifugation (SI Figure 6A) under non-dissociating conditions (using 15 mM MgCl_2_).

Gradient fractions corresponding to fully induced or RbgA-depleted ribosomal particles were pooled and rRNA was stripped of the ribosomal proteins using TRIzol reagent (Invitrogen) followed by isopropanol precipitation for large rRNA yields or using Direct-zol RNA miniprep kit (Zymo Research) for low RNA quantities. RNA has been resuspended and stored in UltraPure water (Thermo Fisher Scientific), and sample concentration was determined using NanoDrop UV-Vis (Thermo Fisher Scientific).

### RNA MS samples and data acquisition

For identification and placement of modifications, ∼10-20 pmol of each ^14^N and ^15^N rRNA were combined (see SI Figure 1 for the complete list of samples and their isotope composition). Each sample was then heat denatured and subjected to RNase T1 or RNase A (∼25-50 units) treatment in a 5 uL volume of 25 mM ammonium acetate (pH 6.0) at 55°C for 1 h. Samples were directly injected into the LC-MS system for MS or MS/MS data acquisition.

Oligonucleotide data were acquired via Agilent Q-TOF 6520-ESI system using negative ion detection following previously published protocol (28). Briefly, nucleolytic oligonucleotides were separated on XBridge C18 column (3.5 µm, 1×150 mm, Waters) using 15 mM ammonium acetate (AA, pH 8.8) in water as mobile phase A and 15 mM AA (pH 8.8) in 50% acetonitrile as B, by the linear ramp of 1-15% mobile phase B in 40 min. MS^2^ collision energies were optimized by direct infusion of RNA oligonucleotide standards (28). Targeted MS/MS was pursued to increase the quality of acquired MS^2^ spectra, used for modifications placement. rRNA isolated from RbgA depleted ribosomes were analyzed via data-dependent MS/MS acquisition, and the following precursor ion selection rules were applied: absolute intensity threshold set to 2000 counts, ions with charge of 1 were excluded, and 0.35 min dynamic exclusion window applied after three precursor selection events.

### CMC chemical labeling

Chemical labeling with pseudouridine specific CMCT (*N*-cyclohexyl-*N*′-(2-morpholinoethyl)carbodiimide metho-p-toluenesulfonate (Sigma-Aldrich) reagent was pursued to identify and quantify pseudouridine modifications in *B. subtilis* rRNA. For that, an equimolar mixture of ^14^N and ^15^N 16S or 23S RNA was freeze-dried, and the obtained RNA pellet dissolved in 20 μl volume of freshly made 0.4 M CMCT in BEU buffer (50 mM Bicine pH 8.5, 4 mM EDTA pH 8.0, 7 M urea). The mixture was incubated at 37°C for 1 h. Then, 480 µL of RNase-free water was added and the excess of unreacted CMCT was removed by two consecutive rounds of Amicon Ultra (0.5 ml, 30 kDa) filtration. To release the label from CMC-guanosines and CMC-uridine adducts present following the reaction, an equal volume (∼ 40 μl) of 100 mM sodium carbonate-bicarbonate buffer (pH 10.5) was added to CMCT free rRNA and incubated at 50°C for 2 h. After two rounds of 30 kDa Amicon Ultra buffer exchange to RNase-free water, derivatized rRNA was dried in a speed-vac system and subjected to T1 or A digestion. LC-MS chromatography gradient was modified to accommodate increased retention of CMC-labeled oligonucleotides on the C18 column and included step 1: 1% of mobile phase B for 5 min; step 2: 1-15% of B in 45 min; step 3: 15-35% of B in 10 min; step 4: 35-100% of B in 5 min.

### Survey of RNA modifications using stable-isotope labeling and least-squares fitting of MS^1^ spectra

The initial phase of the modification search employed an in-house software suit Massacre for mass spectra extraction, m/z matching to RNA digested *in-silico*, and MS envelope fitting using the previously published algorithm Isodist (41). Raw MS^1^ data were converted to mzML file format via MSConvert (ProteoWizard 3.0) and used as Massacre input. *B. subtilis* strain 168 rrnA-16s and rrnA-23S gene sequences were cleaved *in-silico* according to T1 or A specificities with up to 5 missed cleavages. Modification search was enabled by appending at least one and up to four mass tags to the mass of an unmodified nucleolytic oligo. A maximum of 4 methyl group tags (+14 Da or +17 Da with C[^2^H]_3_-methionine labeling), 2 dihydrouridine tags (+2 Da), and 1 hydroxycytidine tag (+16 Da) per oligonucleotide were allowed at a time during stage 1 of discovery (SI Figure 1), and a maximum of 3 pseudouridine tags (-1 Da for 5,6-[^2^H]-labeling and +251.2 Da for CMCT derivatization) at stage 2. Isotope composition for modified and canonical residues was adjusted for the presence of ^15^N and ^2^H. Each Massacre run was configured to find coeluting peaks pairs that match predicted ^14^N and ^15^N masses of the modified oligos present in the *in-silico* digest within 20 ppm tolerance window. Then, extracted MS^1^ envelopes were fit using the element and isotope composition of nucleolytic RNAs. Obtained fits were first filtered using ^14^N /^15^N peak intensity ratio and fit quality metric, and the remaining ones were visually inspected to pick candidate oligos for targeted MS/MS acquisition and sequence analysis.

### MS^2^ spectra identification using Pytheas database search

Pytheas software was used for annotation of MS^2^ spectra, assignment of oligonucleotide sequence, and sequence placement of the unknown modifications (28). MS^2^ scans acquired via targeted MS/MS acquisition were averaged across the width of the chromatographic peak and then exported to mgf format via MassHunter Qualitative Analysis B.07.00, limiting the number of peaks to 250 most intense. Data were matched across all theoretical spectra in the database generated using Pytheas *in silico* digestion routine. During digestion, *B. subtilis* strain 168 rrnA-16S or rrnA-23S sequences were cut according to the RNase specificity with 1 or 3 missed cleavages, and redundant oligonucleotides were consolidated. Then, a custom script was added to the standard workflow that partially replaces canonical with methylated nucleosides: [mA], [mG], [mC], [mU], [mmC], [mmU], [mmA], [mmG], with dihydrouridine [D] or with 5-hydroxycytidine [ho5C] (see SI Table 2 for notations). A maximum of 3 modifications were allowed per oligo. All the methyl groups were allocated to the nucleobase since base and ribose methylations cause small changes to the theoretical spectrum. 12 and 40 ppm precursor and fragment mass tolerance windows were used for matching, and alpha and beta rewards for consecutive matches were set to 2 and 0.025. After the assignment of methylated nucleosides and dihydrouridine was complete (stage 1, SI Figure 1), a new set of spectral databases was generated for pseudouridine identification (stage 2). Positions of pseudouridines encoded by Pytheas as [Y] (5,6-[^2^H] labeling) or [cmc-Y] (CMC labeling) were varied (up to 3 pseudouridines per oligo allowed) while positions of methylations and dihydrouridine were fixed. Upon completion of the database matching, the best-scored sequences were pulled and used for modification mapping. The Ariadne online calculator tool was routinely used for quick verification of the precursor and fragment m/z values when needed (42).

### Quantitative analysis of modifications in RbgA depleted ribosomal particles

rRNAs isolated from unlabeled 45S^RbgA^ and 70S^RbgA^ were spiked with 23S purified from ^15^N-labeled RB1202 cells (1 mM IPTG induction) in approximately 1:1 molar ratio, then digested with RNase T1 or A. Pseudouridines detection was enabled using CMC chemical labeling. Positions of 23S modifications were fixed and provided in the form of the Pytheas input file. The Pytheas final report, containing a list of identified ^14^N and ^15^N nucleolytic oligos (IDs), and MS^1^ traces in mzML format were passed to Massacre for RNA quantitation. The updated Massacre workflow for RNA included 1) consolidation of multiple IDs with the same sequence, 2) LC peak picking using precursor m/z and the retention time window, 3) extraction and fitting of the MS^1^ envelopes. ^14^N/^15^N intensity ratios from the Massacre fits were normalized to 23S RNA input (median ^14^N/^15^N value obtained for 23S IDs that lack modifications) and used to report levels of rRNA modifications.

### Validation of *B. subtilis* genes corresponding to helix 69 modifications in 23S

Deletion strains BKK24260 (*yqxC*::kan), BKK11620(*yjbO*::kan), BKK09210(*yhcT*::kan), BKK15460(*ylyB*::kan) (43) and the reference *B. subtilis* strain 168 were purchased from BGSC (Ohio State University). MGNA-B814 (*yydA*::erm) (44) was obtained from NBRP *B. subtilis* (NIG, Japan). Following initial antibiotic selection, batch cultures were grown in LB medium at 37°C until reached 0.5-0.7 OD_600_. 70S ribosomes were pelleted through the 40% (w/v) sucrose cushion. rRNA, deproteinated using Trizol reagent, was first subjected to treatment with the CMCT reagent, then digested with either RNase T1 or A. Digestion products were analyzed using data-dependent MS/MS acquisition followed by the Pytheas search to identify rRNA modifications affected by the deletion of a single gene. T1 and A datasets are available for download from the PRIDE archive (PXD051518).

### Double cleavage RNase H assay

2’-deoxy-2’-O-methyl chimera oligonucleotides targeting specific sites in 16S and 23S (SI Table 4) were mixed in 2:1 molar ratio with 25 pmol of rRNA in 5 µL volume of the annealing buffer (10 mM Tris-HCl, pH 7.8, 50 mM NaCl, 0.5 mM EDTA) and boiled at 95°C for 2 min. After cooling down to the ambient temperature, 1 µL of 0.1 M DTT and 1 µL of the 10X reaction buffer (500 mM NaCl, 50 mM MgCl_2_, 450 mM Tris-HCl, pH 7.8) were added to the annealed RNAs. 2-3 U of *E. coli* RNase H (New England Biolabs) was then used to initiate a sequence-specific rRNA cleavage in 10 µL final volume, and the reaction was incubated at 37°C overnight. Upon completion, salts and buffer components of the reaction were depleted using Amicon filtration (0.5 ml, 3 kDa), and the RNA retained by the membrane dried in the speed-vac. The resulting double cleavage 5’-P and 3’-OH products were analyzed using data-dependent MS/MS, and positions of modifications (or their absence) were inferred using Pytheas search.

### Ribonucleoside analysis

Complete hydrolysis of rRNA was performed according to the protocol published by the Kellner Lab (45). For relative quantitative analysis of nucleosides, ∼ 20 pmol of each *B. subtilis* and *E. coli r*RNA (isolated from BW25113 strain, grown in M9 minimal medium) with distinct isotope composition between the two species were mixed. RNA digestion was carried out in the final volume of 20 µL by adding 4 µL of 5x digestion buffer (100 mM Tris-HCl, pH 8.0 and 5 mM MgCl_2_), 2 U of Benzonase (Novagen), 0.2 U of Phosphodiesterase I (Worthington), 2 U of quick CIP (NEB), and water as needed. After 2 h incubation at 37°C, 480 uL of RNase-free water was added to the hydrolyzed RNA and proteins were removed by ultrafiltration using Amicon Ultra (0.5 ml, 10 kDa) filter. The filtrate was dried and after resuspension in LC-MS water, the material was analyzed by Agilent Q-TOF 6520-ESI.

To confirm correct nucleoside identification, samples containing ^15^N-labeled *B. subtilis* RNA were prepared and spiked with a mixture of commercially available ribonucleoside standards (SI Table 4). To disambiguate between positional isomers (e.g., 7-methylguanosine, N2-methylguanosine, 2’-O-methylguanosine in 23S), they were pre-mixed at different stoichiometric ratios. Nucleoside retention times and elution order were recorded and used for their subsequent identification in bacteria samples devoid of synthetic standards.

Content of the rRNA hydrolysate was resolved using Zorbax SB-C18 column (5 µm, 0.5 x 150 mm, Agilent) and the LC conditions were tuned empirically, mostly to optimize the detection of modified pyrimidines. Two sets of conditions were established, using either mobile phases A1 (0.1% formic acid in water) and B1(0.1% formic acid in acetonitrile) at column temperature of 30°C, or mobile phases A2 (10 mM ammonium acetate pH 5.4 in water) and B2 (80% acetonitrile in water) without additional column heating. The LC gradient consisted of step 1: 3% of mobile phase B1 (or B2) for 3.5 min; step 2: 3-30% of B1 (or B2) in 6.5 min; and step 3: 30-90% of B1 (or B2) in 5 min. MS detection was carried out using positive ionization in 160-700 m/z range with 1 s MS^1^ scan rate. The capillary voltage was set to 4.5 kV and fragmentor to 100 V. Each biological replicate sample was analyzed twice, injecting an equivalent of 0.2-0.5 pmol of *B. subtilis* rRNA to accurately measure canonical ribonucleosides, used for calculating rRNA input. Furthermore, 5-10 pmol of rRNA was injected for relative quantitation of modified nucleosides, present at quantities that are ∼100-800 fold lower. ^14^N- and ^15^N-peak areas were obtained using integration of extracted ionic chromatograms implemented in MassHunter Qualitative Analysis B.07.00. For *B. subtilis* labeled with 5,6-[^2^H]-uracil, MS^1^ data were fitted, and ^14^N- and ^15^N-peak intensities calculated using Massacre. To correct the data for rRNA input, *B. subtilis* / *E. coli* peak area ratio for each modified nucleosides was normalized to the *B. subtilis / E. coli* peak area ratio for the canonical nucleosides (i.e., median value for A, G, U, C) and reported in Figure 3.

**Table 1.**
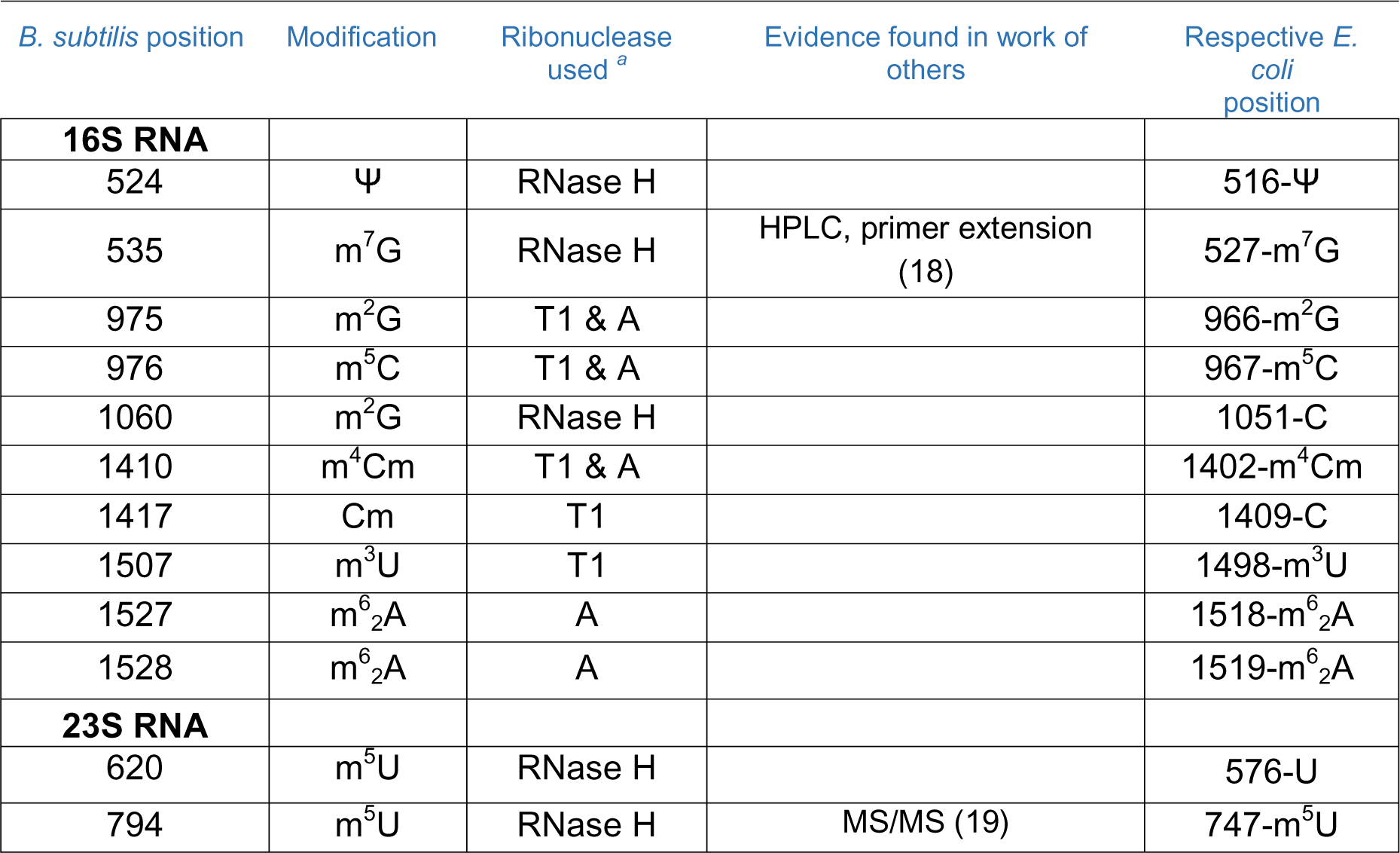
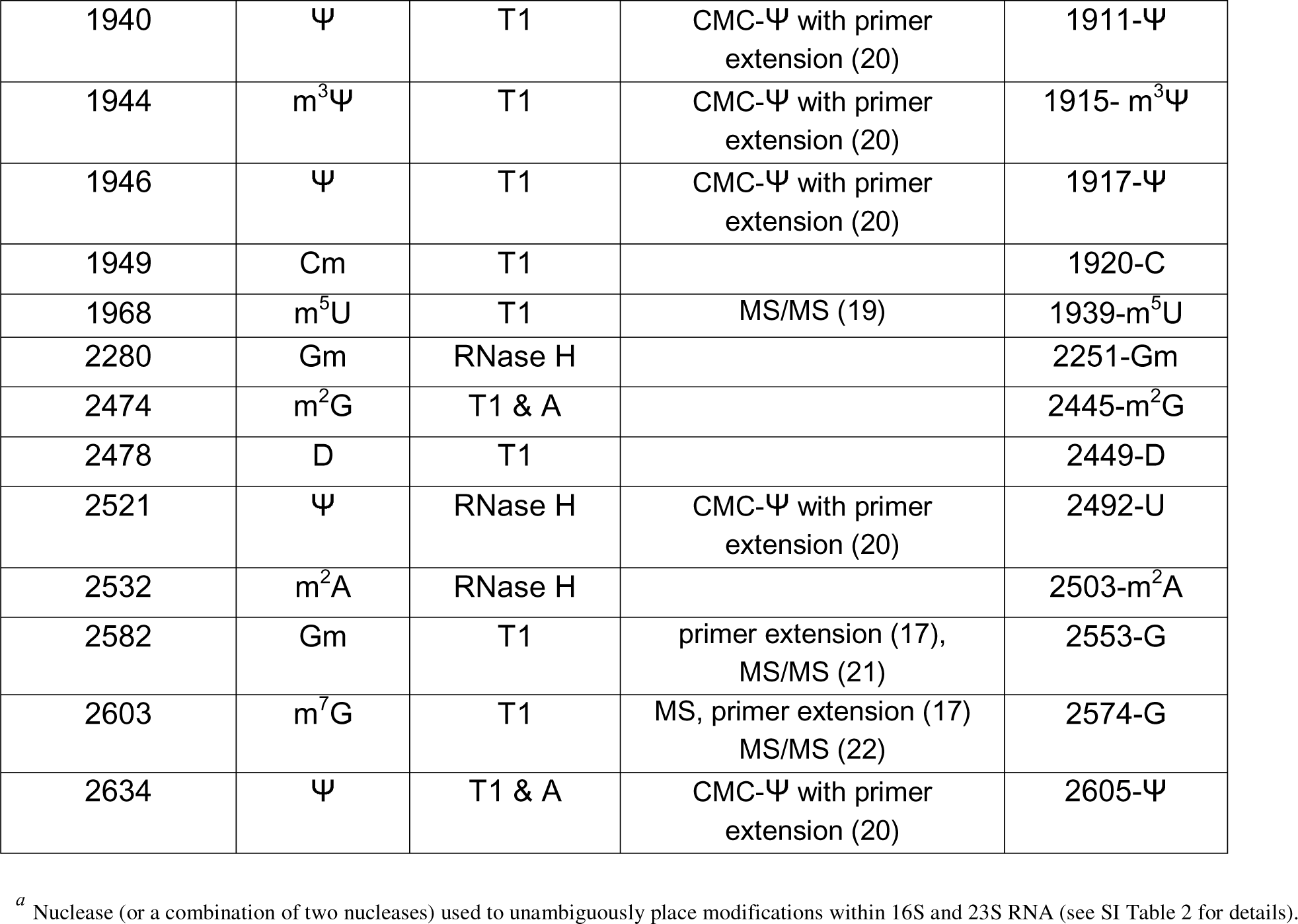
RNA modifications identified in *B. subtilis* 16S and 23S RNA.

| *B. subtilis* position | Modification | Ribonuclease used <sup>a</sup> | Evidence found in work of others | Respective *E. coli* position |
| --- | --- | --- | --- | --- |
| **16S RNA** |  |  |  |  |
| 524 | $\Psi$ | RNase H | | 516-Ψ |
| 535 | m<sup>7</sup>G | RNase H | HPLC, primer extension (18) | 527-m<sup>7</sup>G |
| 975 | m<sup>2</sup>G | T1 & A |  | 966-m<sup>2</sup>G |
| 976 | m<sup>5</sup>C | T1 & A |  | 967-m<sup>5</sup>C |
| 1060 | m<sup>2</sup>G | RNase H |  | 1051-C |
| 1410 | m<sup>4</sup>Cm | T1 & A |  | 1402-m<sup>4</sup>Cm |
| 1417 | Cm | T1 |  | 1409-C |
| 1507 | m<sup>3</sup>U | T1 |  | 1498-m<sup>3</sup>U |
| 1527 | m<sup>6</sup><sub>2</sub>A | A |  | 1518-m<sup>6</sup><sub>2</sub>A |
| 1528 | m<sup>6</sup><sub>2</sub>A | A |  | 1519-m<sup>6</sup><sub>2</sub>A |
| **23S RNA** |  |  |  |  |
| 620 | m<sup>5</sup>U | RNase H |  | 576-U |
| 794 | m<sup>5</sup>U | RNase H | MS/MS (19) | 747-m<sup>5</sup>U |
| 1940 | $\Psi$ | T1 | CMC-Ψ with primer extension (20) | 1911-Ψ |
| 1944 | $m^3\Psi$ | T1 | CMC-Ψ with primer extension (20) | 1915- $m^3\Psi$ |
| 1946 | $\Psi$ | T1 | CMC-Ψ with primer extension (20) | 1917-Ψ |
| 1949 | Cm | T1 |  | 1920-C |
| 1968 | $m^5U$ | T1 | MS/MS (19) | 1939-$m^5U$ |
| 2280 | Gm | RNase H |  | 2251-Gm |
| 2474 | $m^2G$ | T1 & A | | 2445-$m^2G$ |
| 2478 | D | T1 |  | 2449-D |
| 2521 | $\Psi$ | RNase H | CMC-Ψ with primer extension (20) | 2492-U |
| 2532 | $m^2A$ | RNase H | | 2503-$m^2A$ |
| 2582 | Gm | T1 | primer extension (17), MS/MS (21) | 2553-G |
| 2603 | $m^7G$ | T1 | MS, primer extension (17) MS/MS (22) | 2574-G |
| 2634 | $\Psi$ | T1 & A | CMC-Ψ with primer extension (20) | 2605-Ψ |

<sup>a</sup> Nuclease (or a combination of two nucleases) used to unambiguously place modifications within 16S and 23S RNA (see SI Table 2 for details).

To enhance ionization efficiency and avoid interference from coeluting species, several uridine derivatives (m^3^Ψ, Ψ, D) were interrogated using negative detection mode and mobile phases A2 and B2. Coelution (3.0 min column retention) and small (+2 Da) mass shift between Ψ and D, preclude their independent measurements in 23S RNA using MS^1^ signal alone (SI Figure 5B). Therefore, Ψ nucleoside quantification was carried out via two MRM transitions 243.06→153.03 and 243.06→183.05 for ^14^N precursor, and 245.06 →155.02 and 245.06 →185.04 for ^15^N at a collision energy of 10 V. 5,6-dihydrouridine quantification was not attempted.

## RESULTS

### The overall strategy for discovery of RNA modifications

The standard bottom-up approach has been utilized for the discovery of modifications, where 16S and 23S RNA were purified from ^14^N and ^15^N labeled *B. subtilis* cells and subjected to either RNase T1 or A treatment. The resulting mixture of nucleolytic oligos (∼ 3-15 nt size range) was first analyzed using LC-MS. Then precursor ions containing specific MS^1^ spectral signatures were interrogated by tandem MS. Given that base/ribose methylations and pseudouridines are the most abundant rRNA modifications (SI Table 1) in prokaryotes, the discovery workflow has been split into two parts: discovery of methylations and other non-mass silent modifications (5-hydroxycytidine and 5,6-dihydrouridine were included in the search), and the discovery of pseudouridines (see stages 1 and 2 in SI Figure 1). To amplify spectral features of the methylated oligonucleotides and to discriminate pseudouridines from isomeric uridines, specific isotope and chemical labeling schemes were employed at the sample preparation step.

During the precursor (i.e., MS^1^) survey phase of the discovery workflow, subsets of candidate sequences were identified using predictable mass shifts and isotopic envelope transformations induced by known RNA modifications. Briefly, monoisotopic masses of the coeluting ^14^N and ^15^N peak pairs were extracted from the raw data and matched to the masses of oligos present in the theoretical digest, then experimental ^14^N and ^15^N spectra were fit using the provided atomic composition (solid line fits in Figure 1A and 1C). This step efficiently eliminated canonical sequences, leaving a small number of “mass-shifted” precursors (∼15) for targeted LC-MS/MS. Targeted acquisition notably improved the quality of the tandem spectra enabling sequence assignment and localization of the modified nucleoside with high confidence. Pseudouridine modifications (Ψ) are isobaric to uridines, and reliable detection requires specific metabolic labeling or chemical treatment for effective “mass-shifting” of Ψ in RNA. Consequently, pseudouridine search was carried out independently (at stage 2), after the localization of other modifications was complete, following the exact same data acquisition and analysis scheme.

**Figure 1.**
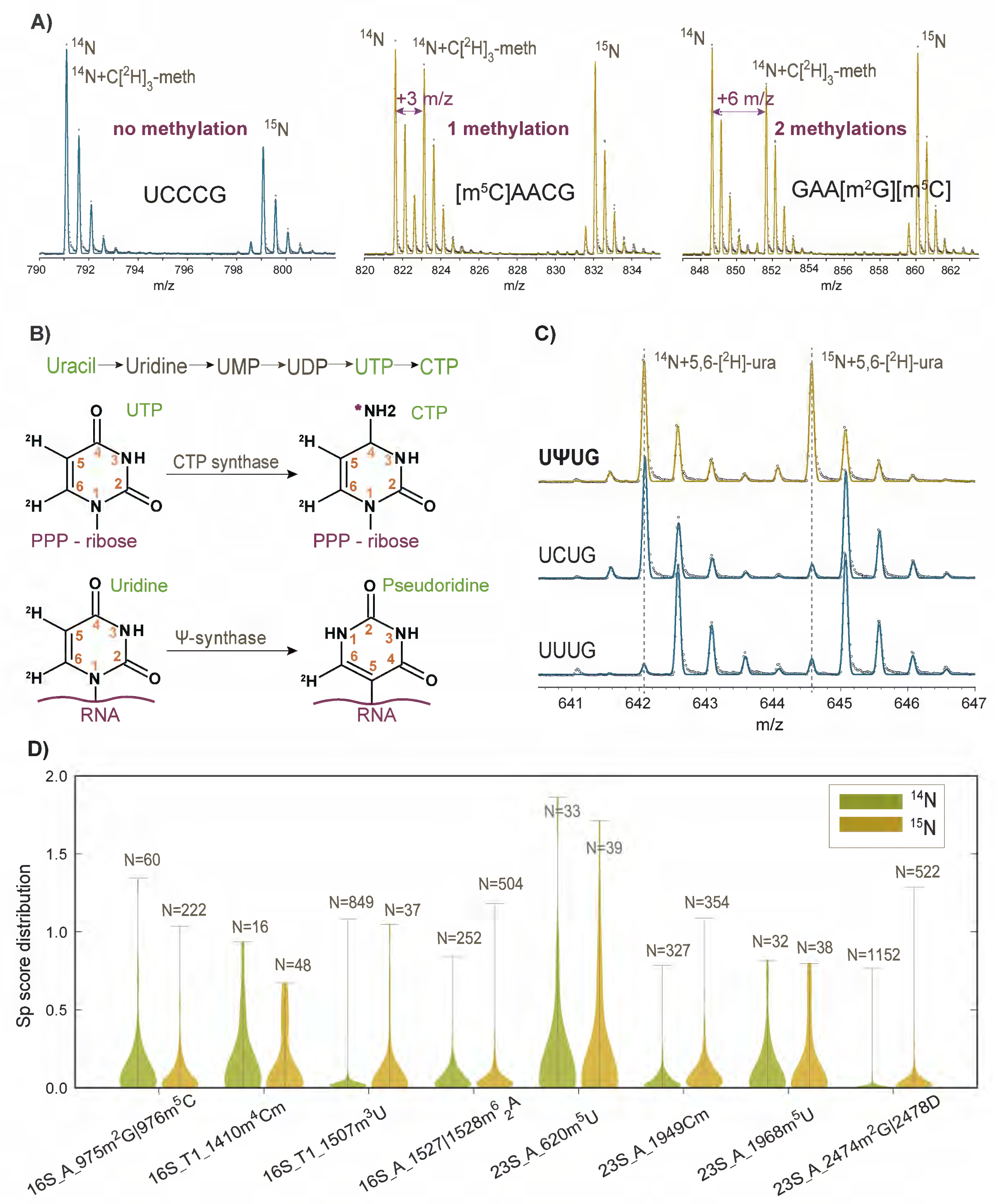
Stable isotope labeling, quantitative MS^1^, and oligonucleotide MS^2^ tools for identification of modifications in *B. subtilis* rRNA. (A) Examples from MS^1^ datasets employing ∼1:1:1 stoichiometric mixture of three isotopically distinct 16S species: ^14^N, ^14^N +C[^2^H]_3_-methionine, and ^15^N labeled. Nucleolytic oligos without a methyl group observe two population spectra with 2:1 intensity ratio, while methylations result in three population spectra with 1:1:1 ratio. Note, that the latter ratio can slightly vary due to increased retention of deuterated RNA on the C18 column. Single methyl group gives rise to +3 Da (or +3 m/z as shown) mass shift between ^14^N and ^14^N +C[^2^H]_3_-methionine peaks, two methyl groups to +6 Da (+6 m/z) shift etc. (B) Schematic of the pyrimidine [^2^H]-labeling in *B. subtilis* culture using 5,6-[^2^H]-uracil as a metabolic precursor. The purple star marks the nitrogen atom in CTP that is ^15^N-labeled when 5,6-[^2^H]-uracil is added to the ^15^N-substituted minimal medium. (C) MS^1^ data demonstrates the utility of the labeling scheme (B) to resolve pseudouridines from canonical uridine and cytidine nucleosides. 4-mer oligos spectra, identified in T1 digested 23S, are different in a single pyrimidine nucleoside. Characteristic 1 Da (or 1 m/z as shown) shifts are observed between ^14^N-species (Ψ, C vs U) and ^15^N-species (Ψ vs C, U), permitting identification of all three. In (A) and (C), MS fits (solid line) generated using Isodist are overlayed on top of the raw data (grey dots). (D) Pytheas S_p_ score distributions for selected ^14^N and ^15^N precursor ions using MS^2^ variable modifications search. The number of competing oligonucleotide spectrum matches (N) is shown, and the top horizontal bar indicates S_p_ value for the top sequence match. For each entry, RNA sample (16S or 23S), ribonuclease (T1 or A), position, and identity of the modification are shown. Data for ^14^N and ^15^N precursor pairs are in green and in yellow. Detailed Pytheas outputs used for plotting are provided in Supplementary Data.

Elicited mass changes to precursor and fragment ions were accommodated by specifying element and isotope composition of the modified nucleosides during *in-silico* digestion step. Variable modifications (modifications that are either present or absent) were sampled explicitly, in all possible arrangements across the sequence space provided by 16S or 23S RNA cleaved by the RNase. The upper limit to the net number of modifications per nucleolytic oligo was set to 3-4 (see Material and Methods for details), which generated a search space of manageable complexity (SI Table 3) but allowed for the presence of closely spaced modification clusters in rRNA.

Using the described workflow, we detected 15 new and 10 previously identified *B. subtilis* modifications (Table 1). Each assignment was confirmed by multiple means: using different enzymatic cleavage products, by varying isotope composition of rRNA, and in the case of pseudouridines by using chemical tagging. Modifications type and positions presented here are fully consistent with prior experimental evidence collected by other labs using HPLC and low-resolution techniques such as sequencing gels (17,18,20), and agree well with modifications that were annotated in *E. coli* and other bacteria (SI Table 1). Excitingly, this further broadens the landscape of functional modification sites in RNA biology.

### Isotope and chemical labeling for identification of methylations and pseudouridines

To identify a list of digestion products that potentially carry one or many methyl groups, we grew cells in C[^2^H]_3_-methionine supplemented minimal medium (^14^N-C[^2^H]_3_-meth, Figure 1A). C[^2^H]_3_-methionine is a metabolic precursor of SAM, a co-factor used by cell methyltransferases to catalyze the transfer of C[^2^H]_3_ moiety to RNA. Consequently, a characteristic +3 Da mass shift becomes a prominent MS signature of the methyl group presence. In samples containing a nearly equimolar mixture of ^14^N-, ^15^N-, and ^14^N-C[^2^H]_3_-methionine labeled RNAs, unmethylated oligos present two peak spectra with a 2:1 intensity ratio (UCCCG in Figure 1A, note merging of ^14^N states). Methylated oligos, on the other hand, exhibit three peak spectra with an approximately 1:1:1 ratio ([m^5^C]AACG and GAA[m^2^G][m^5^C]). The +3 Da shift per methyl group rule applied to all methylated precursors detected in this study except for T1 and A cleaved fragments subsequently assigned to m^2^A at position 2532 in 23S. 2532-m^2^A produces +2 Da mass shift and is equivalent to the RlmN dependent 2503-m^2^A methylation in *E. coli* (Table 1). RlmN belongs to the family of methyltransferases with a distinct radical-based mechanism, where two participating SAM molecules result in a bulk transfer of C[^2^H]_2_ rather than C[^2^H]_3_, consistent with the observed +2 Da mass shift (46). This observation is in line with the broad conservation of RlmN across the bacteria phyla (47) and in *B. subtilis* (SI Figure 2B).

Two complementary approaches were undertaken to make the normally isobaric rRNA pseudouridines apparent via MS measurements, one using metabolic labeling in cell culture, and the other exploiting the carbodiimide reactivity of pseudouridines (SI Figure 1). The first approach relies on the efficient incorporation of the deuterium atom into C5 of bacteria uridines, as demonstrated in prior work (35). If 5,6-[^2^H]-uracil precursor is available during bacteria growth, it is efficiently assimilated into 5,6-[^2^H]-uridine and 5,6-[^2^H]-cytidine triphosphates via the salvage pathway of pyrimidine biosynthesis (Figure 1B). After rRNA is transcribed, pseudouridine synthases carrying out uridine isomerization, exchange C5-^2^H to C5-^1^H leading to the detectable -1 Da shift between U and Ψ. An example in Figure 1C shows ^14^N+5.6-[^2^H]-ura and ^15^N+5.6-[^2^H]-ura spectra pairs for three oligonucleotides identified in T1 treated 23S following the metabolic labeling. Clearly, the difference in a single pyrimidine residue (Ψ vs C vs U, from top to bottom) gives rise to distinct -1 Da mass shifts between U and Ψ ^14^N spectra and between C and Ψ ^15^N spectra, permitting recognition of Ψ containing precursors. Pyrimidine-specific 1 Da shifts shown here using MS^1^ data are further retained in the fragmentation spectra of ^14^N and ^15^N species that are directly available for MS^2^ based oligo sequencing to ultimately confirm or identify new positions of Ψ modifications. This has been done previously for pseudouridine detection in yeast rRNA (28) and is fully exploited here (SI Table 2).

The alternative route used for pseudouridine profiling is a chemical reaction with a bulky CMC (N-cyclohexyl-N′-(2-morpholinoethyl)carbodiimide) group that specifically tags pseudouridines within a canonical RNA sequence. Taking advantage of CMC-Ψ specific stalling of the reverse transcriptase (RT), many labs combined RT primer extension and gel-sequencing, later replaced by more robust NGS technologies to *de novo* identify pseudouridines in cellular RNAs including bacteria rRNA (20,27). CMC derivatization has been adapted for use in mass spectrometry by the Limbach group (48,49). CMC adds a characteristic 251.2 Da mass tag per single pseudouridine, causes an RNase A missed cleavage at the CMC-Ψ site (SI Table 2), and dramatically increases retention of the derivatized product on the reverse-phase column, making CMC-tagged oligos easy to identify out of hundreds of unlabeled and often coeluting RNase digestion products. These advantages were exploited here to find pseudouridine precursor ions and characterize their sequence via MS/MS (SI Figure 3).

### MS^2^ spectra analysis reveals positions of RNA modifications

Quality annotation of an entire MS^2^ spectrum remains critical for reliable sequence determination and modification placement. Following MS^1^-based selection of “mass-shifted” precursor ions, targeted MS/MS acquisition was pursued, and acquired MS^2^ spectra were matched against theoretical databases of spectra using Pytheas software (SI Figure 1). A top-ranking sequence match was accepted as a true ID if confirmed using both ^14^N and ^15^N precursors from at least two replicate experiments. Sequences assigned to the modified oligos from *B. subtilis* samples are listed in SI Table 2 and top scoring spectra are available for download from the PRIDE database (PXD050367).

Unsurprisingly, variable modifications included in the search expand the size of spectral databases, setting high competition between the candidate sequences, with up to 3-4.5·10^5^ sequences present in 16S and 23S theoretical digests (SI Table 3). For each precursor, the number of alternative sequence IDs varied dramatically depending on the rRNA, nuclease used, and isotope composition (Figure 1D and Supplementary Data). For example, RNase A product 616-AGAA[mU]-620 labeled with ^14^N was the top ranking match among 33 other candidate sequences, found in 23S *in-silico* library within 5 ppm of the precursor m/z. Among candidates are AGAAU plus three other anagrams of this sequence with every possible allocation of the methyl group, six unmodified oligos, and seven oligos that contain other modifications. This emphasizes that automated search such as implemented here is the only practical way of managing complexity enumerated using flexible modifications, as it would be hard to verify all the matches by hand.

For the majority of precursors, the quality of acquired MS^2^ spectra was sufficiently high to discriminate between first- and second-best sequence matches, with the median value of the Pytheas relative distance score (ΔS_p_2) equal to 0.43 (calculated for data presented in SI Table 2). Except for a few cases when low ΔS_p_2 values were obtained (see SI Table 2 discussion), Pytheas search could effectively resolve oligonucleotide sequences and localize modification sites.

### Double cleavage RNase H assay for unambiguous placement of modifications within rRNA

The use of commercial ribonucleases T1 (G-specific) and A (U, C-specific) for post-transcriptional modifications mapping is very common but limited due to frequent occurrence of cleavage sites generating short oligonucleotides (1-5 nt) that are hard to uniquely map on to 1.5 kb (16S) and 2.9 kb (23S) rRNA. Long RNAs in general tend to produce short modified T1/A digestion products that can map to multiple sequence sites, and placement of modifications becomes challenging (e.g. [m^2^A]UG and G[m^2^A]U assigned to 2532-m^2^A of 23S, SI Table 2). In total, eight modification sites in 16S and 23S could not be uniquely mapped using either T1 or A alone or assuming true overlap between T1 and A cleavage products (Table 1, marked by RNase H). While newly discovered enzymes and cleavage tools (29,30) that potentially increase oligo size remain unavailable to the broad public, we found a new way to repurpose RNase H enzyme for the needs of oligonucleotide mass spectrometry.

RNase H cleavage method used here was adapted from the work of Zhao and Yu (50) who proposed the design of synthetic 2’-deoxy-2’-O-methyl complementary sequences to cut RNA in a precise sequence-specific manner. For *B. subtilis* rRNA, similar chimera probes that contain two cleavage sites separated by the 7-9 nt long region complementary to the rRNA area of interest were used (Figure 2A). The new assay was very effective, producing a dominant MS^1^ peak corresponding to the anticipated 3’-P 5’-OH double cleavage product with small quantities of the 3’-dephosphorylated (3’-OH 5’-OH) byproduct. No significant off-target cleavages were detected, however some of the targets had an elevated propensity for ± 1 nt slipped cleavages, thus increasing or decreasing the size of the expected product by 1 nt. To take advantage of this new method, we constructed several RNase H chimera probes against positions that either could not be validated via conventional T1/ A treatment, or against sites suggested by the precursor survey data, and the sites modified in other bacteria species (SI Tables 4 and 5). MS^2^ spectra were collected and Pytheas search carried out to map modifications on to rRNA (Figure 2B) or otherwise prove absence of modifications.

**Figure 2.**
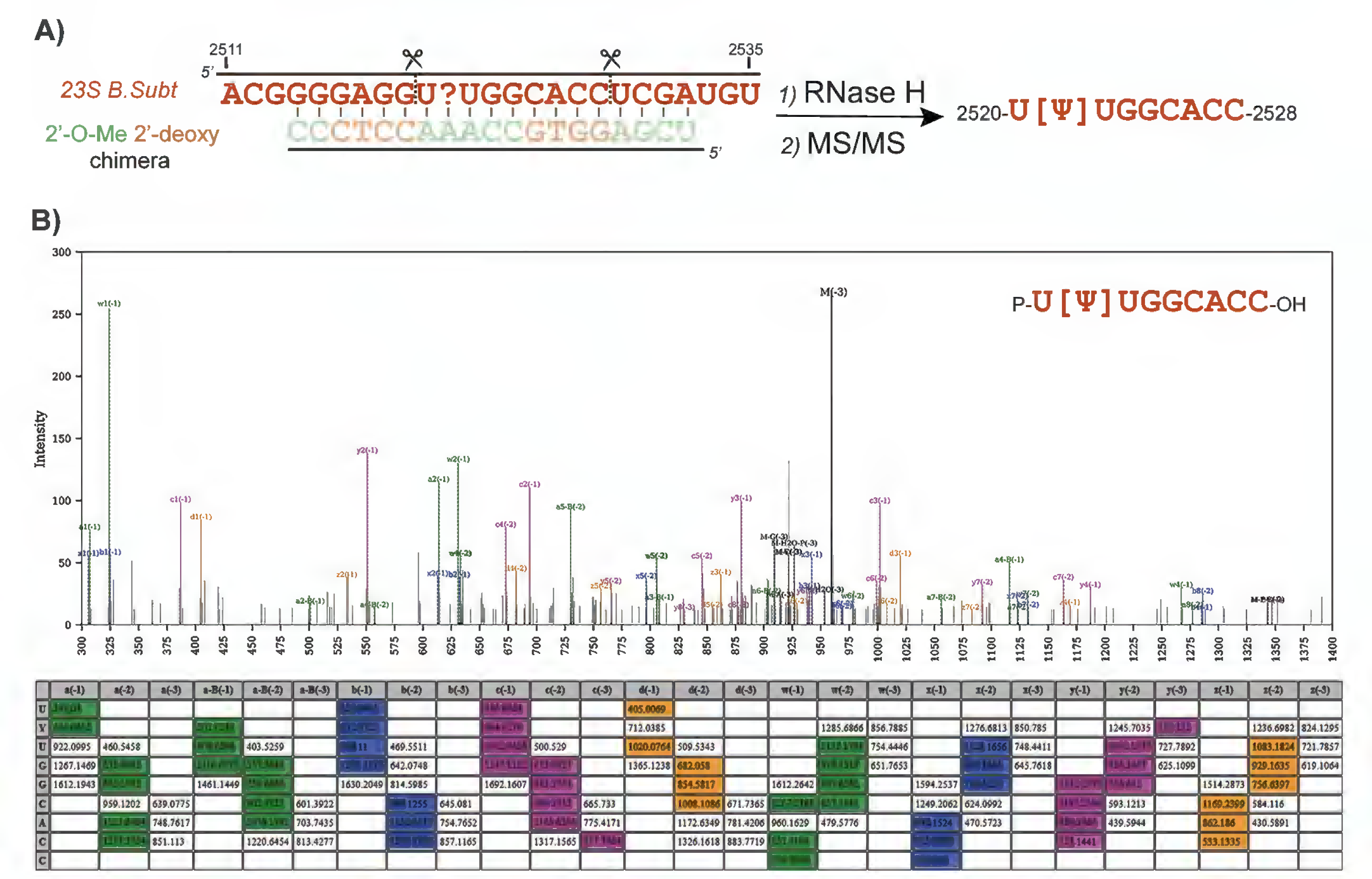
Double cleavage RNase H assay for modification placement. (A) Schematic of the 2’-deoxy-2’-O-methyl chimera design and sequence-guided RNase H cleavage at the interrogated site, followed by mass spectrometry detection (B) Pytheas annotation of the MS^2^ spectrum for the dominant RNase H cleavage product obtained using 23S labeled with 5,6-[^2^H]-uracil. Sequence-directed cleavage method and database search were used to prove pseudouridine modification at position 2521 of *B. subtilis* 23S, which otherwise hard to resolve using conventional nucleolysis. In the spectrum, assigned peaks are colored according to an RNA fragmentation ion series (a/a-B/w – green, b/x – blue , c/y – pink, d/z – yellow). The table below shows m/z values for all predicted MS^2^ fragments, and fragments observed experimentally are highlighted using the color scheme described.

**Figure 3.**
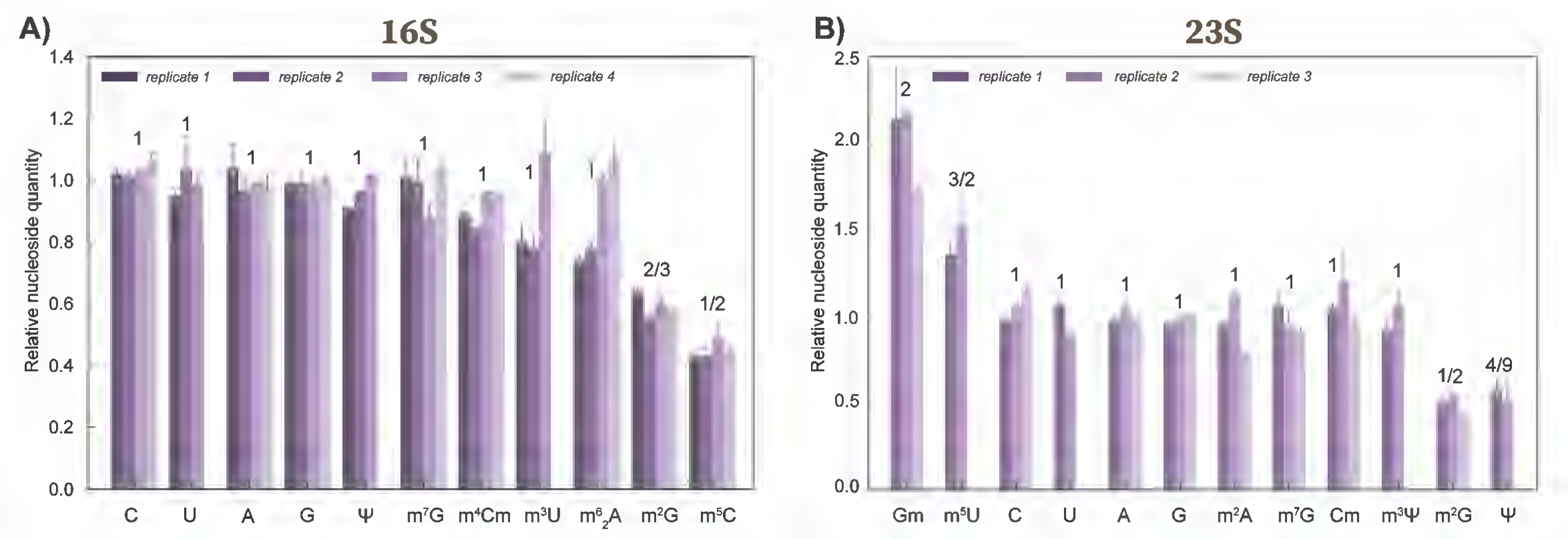
Characterization of *B. subtilis* nucleoside composition via complete 16S and 23S digestion using *E. coli* as a quantitative reference. Each bar represents data from an independent biological replicate of *B. subtilis*. Error bars refer to the standard deviation between multiple technical replicates when available. Numeric labels on top of each bar indicate the expected nucleoside ratio between two species, assuming that each site is 100% modified. For example, an expected ratio equal to 2 for Gm and ½ for m^2^G in (B) indicates that *B. subtilis* 23S RNA should have twice as many Gm and twice as low m^2^G residues as *E. coli*. Absolute quantities of each nucleoside in *B. subtilis* RNA were calculated based on the bottom-up MS inventory (Table 1). In (A), replicates 1 and 3 used ^14^N-16S of *B. subtilis,* replicate 2 – ^15^N-16S, and replicate 4 – ^15^N+5.6- [^2^H]-uracil-16S. In (B), replicate 1 used ^15^N-23S, replicate 2 – ^14^N-23S, and replicate 3 – ^15^N+5.6-[^2^H]-uracil-23S. *E. coli* rRNA of complementary isotope composition was used as an internal reference.

While RNase H double-cleavage method played a decisive role to complete placement of modifications on rRNA sequence, it will be broadly applicable for MS-based interrogation of specific sites in virtually any RNA of synthetic or biological origin.

### Nucleoside content of 16S and 23S

Bacterial rRNA and tRNA carry a large yet manageable repertoire of methylated nucleosides, spanning from ribose methylations to different positional isomers of the methylated base, that is largely determined by the evolutionary conservation of methyltransferases in prokaryotes (47). Despite being tricky to resolve, we found that oligonucleotide MS data carry information that can help to distinguish positional isomers, or at least to narrow down the number of possible methylation types.

It is recognized that modifications, including positional isomers, influence column retention, therefore synthetic standards can be used to verify modification chemistry. While the availability of modified oligonucleotides is limited, many labs use biological RNAs as a reference. Given an overall high conservation of rRNA sequence between species and complete annotation of *E. coli* modifications, a sample containing ^14^N-RNA from *B. subtilis* was spiked with ^15^N-RNA from *E coli*. The analysis revealed eight coeluting ^14^N/^15^N peak pairs (SI Table 6), strongly suggesting that sites found are where both sequence and modification type are identical.

Furthermore, RNases T1 and A are well known to create missed cleavages at certain modified nucleosides: T1 misses Gm, m^7^G, and m^1^G, but cleaves at m^2^G, while A misses Um, Cm, m^3^U, m^3^Ψ and cleaves at m^5^U and m^5^C. Besides, MS^2^ spectra contain evidence of the specific modification type such as m^7^G due to their unique fragmentation behavior (SI Figure 4C) or otherwise display fragments corresponding to free methylated bases and methylated base losses (SI Table 2). Spectra with multiple methylations carry features that help to attribute both methyl groups to a single base ([mmA] base loss corresponding to 1527|1528-m^6^_2_A, SI Figure 4A), or to assign methylations to adjacent positions ([mG] and [mC] base losses corresponding to 975-m^2^G and 976-m^5^C in SI Figure 4B). MS^2^ fragments containing m^5^U, m^5^C, or any methylated pseudouridine (m^3^Ψ, m^1^ Ψ or Ψm) observe a characteristic -1 Da mass shifts when rRNA is labeled with 5.6-[^2^H]-uracil. m^3^Ψ is easily confused with m^5^U, as neither of the two react with CMCT, and both are found in bacterial 23S. Unlike m^5^U, m^3^Ψ is unreceptive to RNase A cleavage (SI Table 2), a feature used to distinguish these two isomers. In summary, nuclease specificities and often MS^2^ spectra themselves provided enough MS-based signatures for assigning positions of G, U, and C methyl groups. Adenosine methylation types remain irresponsive to the nucleolytic treatment used which was addressed using complete rRNA hydrolysis and the nucleoside LC-MS analysis.

A near-equimolar mixture of isotopically labeled *B. subtilis* and *E. coli* rRNA was enzymatically converted to nucleosides, where *E. coli* rRNA served as a quantitative reference with a well-defined nucleoside composition. For 16S, all predicted modification types, except Cm, were identified and quantified, with a good correspondence between expected and measured stoichiometry ratios (Figure 3A and SI Figure 5A). Since *E. coli* 16S lacks Cm, its presence was confirmed separately using ^15^N-16S from *B. subtilis* spiked with a synthetic nucleoside standard. For 23S RNA, coelution of 5,6-dihydrouridine (D) and pseudouridine (Ψ) nucleosides precluded quantitative analysis of D, and Ψ was quantified using multiple reaction monitoring (MRM, SI Figure 5B). Other modified nucleosides, including the only methylated adenosine m^2^A present in 23S were well resolved chromatographically, and their isotope ratios are in good agreement with the expected values. It remains possible that the increased stoichiometry ratio for 23S pseudouridines (∼5/9 vs expected 4/9) is due to one extra Ψ residue that escaped identification at the oligonucleotide level. Otherwise, the result can be explained by the precision of the measurement (namely, MRM transitions chosen for quantification), or by the incomplete pseudouridine stoichiometries in *E. coli* 23S.

To summarize, except for 23S pseudouridines, the total modification content agrees well between the nucleoside analysis and number of modifications detected using bottom-up approach. This completes the identification of rRNA modifications in *B. subtilis*, with a total of 12 types of modified nucleosides found at 25 different positions.

### Inventory of modifications in RbgA-dependent large subunit intermediate

To begin to understand how rRNA modifications are acquired along the biosynthesis pathway we chose to focus on the large subunit (LSU), exploring the key role of the *B. subtilis* GTPase called RbgA during late steps of LSU assembly. Although absent in the *E. coli* genome, *rbgA* (*ylqf*) is an essential gene, necessary for the assembly process to proceed past an RbgA dependent 45S intermediate (45S^RbgA^). Composition of 45S^RbgA^ has been extensively characterized including MS and EM analyses (40), demonstrating a mixed pool of particles (class A and B) with a well-resolved LSU core and either completely missing or a well-defined CP (central protuberance), and a partially defined L7/L12 stalk region (37). Importantly, all the 45S^RbgA^ particles are missing PTC. To link available structural information with presence of RNA modifications, 45S^RbgA^ intermediate has been isolated from cells grown at permissive conditions (5 or 10 µM IPTG induction of RbgA, SI Figure 6A), and its RNA component analyzed relative to the mature, modified reference (Figure 4A).

**Figure 4.**
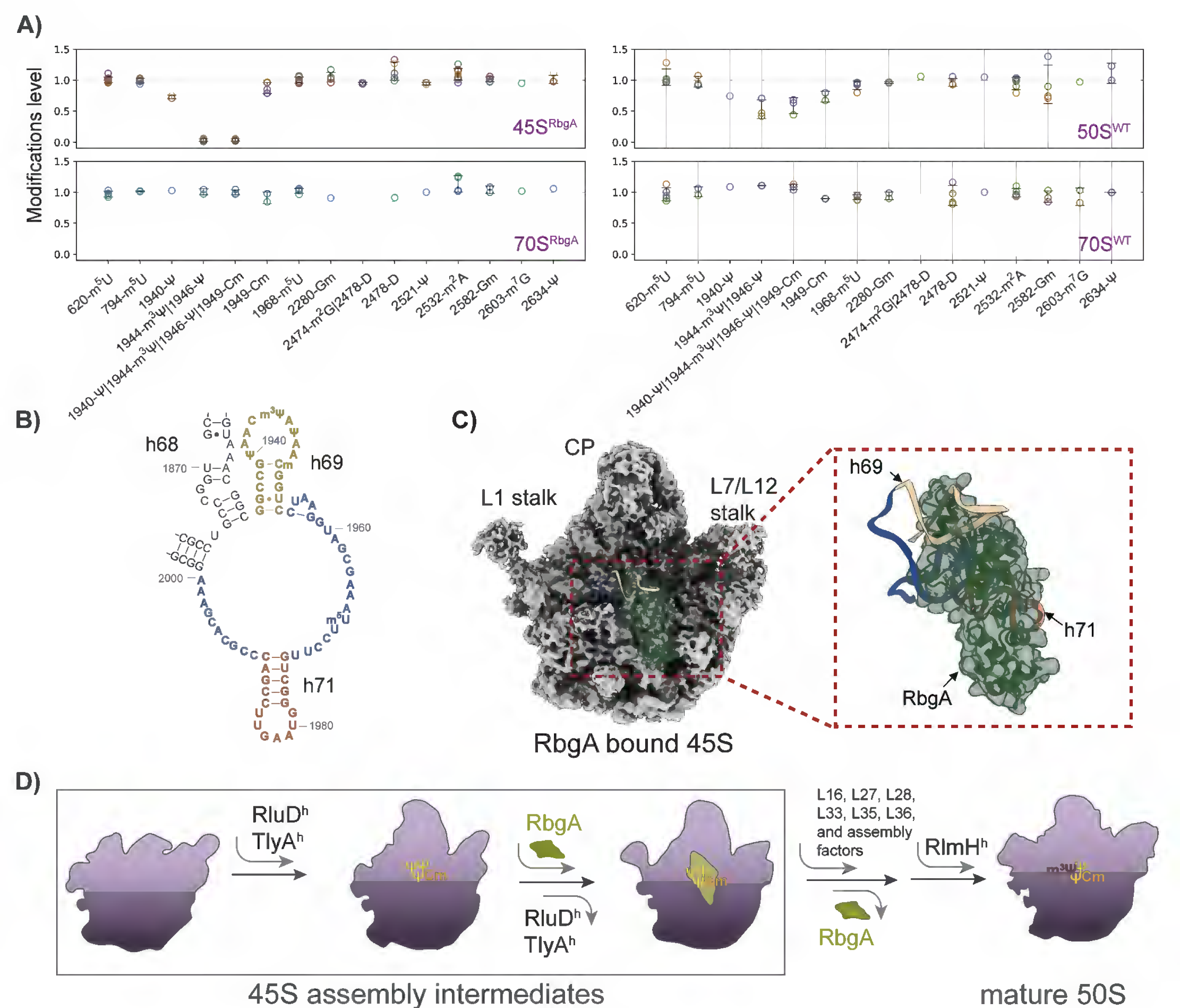
Analysis of rRNA modifications in the late stage of the large subunit assembly. (A) Relative levels of 23S modifications in the ribosomal particles isolated from RbgA-depleted cells □ 45S^RbgA^, 70S^RbgA^ (5 µM and 10 µM IPTG induction, data merged), and from cell with constitutive expression of RbgA (1 mM IPTG) □ 50S^WT^, 70S^WT^. Modifications were quantified relative to ^15^N-23S *B. subtilis* and corrected for the median ^14^N/^15^N ratio. Data for independent biological experiments are shown using various colors, error bars indicate standard deviation for all the datapoints available. Modified residues that are part of a modification cluster can be identified individually and in a group of other modifications (e.g., 1940-Ψ and 1940-Ψ |1944-m^3^Ψ |1946-Ψ |1949-Cm). In such cases, data are shown separately for each combination of modifications. (B) Fragment of 23S secondary structure, indicating positions of the modified residues within h69 and h69-h71 junction. (C) Cryo-EM map of the 45S intermediate bound to the recombinant RbgA (PDB ID: 6PPK), with RbgA shown in green. rRNA helices h69 (yellow), h71 (orange), and h69-h71 junction (blue) are missing in the EM map shown and were modeled from the mature 70S (PDB ID: 3J9W). CP refers to the central protuberance. (D) Schematic of RbgA dependent steps during LSU assembly, integrating late modifications 1940-Ψ, 1944-Ψ, 1946-Ψ by RluD^h^ (YlyB); 1949-Cm by TlyA^h^ (YqxC), and 1944-m^3^Ψ by RlmH^h^ (YydA). These modification enzymes are *E. coli* or *M. tuberculosis* homologs.

Compared to 70S ribosome, modification status of 45S^RbgA^ intermediate is complete except for three pseudouridines and two methylations, all of which reside in helix 69 of 23S (positions 1935-1953, Figure 4B in yellow). A closer look into 45S^RbgA^ data suggests that the pseudouridine methylation 1944-m^3^Ψ is fully missing, but pseudouridines themselves are likely present at high sub-stochiometric ratios (0.73±0.03 calculated for 1940-Ψ, and 1944-Ψ |1946-Ψ spectra presented in SI Figure 6B). Likewise, 50S^WT^ fractions from cells with constitutive expression of RbgA are partially depleted of the three pseudouridines in h69 (0.75 calculated for 1940-Ψ), however 1944-m^3^Ψ is 56±13% present.

The three pseudouridines in the h69 stem loop are universally conserved. *E. coli*, for example, utilizes a single protein enzyme RluD to carry out h69 pseudouridylations, followed by the RlmH dependent methyl transfer to form 1915-m^3^Ψ (corresponds to 1944-m^3^Ψ in *B. subtilis*). Depletion (and 1944-m^3^Ψ absence) of h69 pseudouridine modifications in *B. subtilis* 45S^RbgA^|50S^WT^ is consistent with the late status of RluD in *E. coli* LSU biogenesis (36,38). Moreover, it has been speculated that *E. coli* RlmH may induce its function not until the final steps of the large and the small ribosomal subunits joining (51). Given that a portion of particles associated with the 50S^WT^ peak on the density gradient arise from recycled and dissociated 70S, the partial abundance of 1944-m^3^Ψ in 50S^WT^ is expected. On the other hand, 45S^RbgA^ particles are incomplete assembly intermediates unable to form translation competent ribosomes and thus lack 1944-m^3^Ψ.

Another h69 methylation, 1949-Cm is partially depleted in both 45S^RbgA^ (0.87±0.09) and in 50S^WT^ (0.74±0.09), suggesting that 1949-C methyl transfer might be concurrent to h69 pseudouridines isomerization. *tlyA* dependent ribose methylation at position 1920-Cm (corresponds to 1949-Cm in *B. subtilis*) is found in *M. tuberculosis.* Recently, Laughlin and co-workers demonstrated that recombinant TlyA enzyme stably binds the matured large subunit of *M. tuberculosis* and efficiently catalyzes cytidine methyl transfer (34).

The late status of h69 modifications, suggested by our data, correlates well with missing EM density for helix 69 (as well as h68 and h71) in 45S intermediate particles (PDB IDs: 6PVK, 6PPF) or RbgA bound 45S (6PPK, Figure 4C). In the latter structure, RbgA occupies P site, sterically excluding and preventing docking of h69 and h71 in their final conformation. By doing so, RbgA likely blocks access to the modification enzymes which remain to act either before RbgA binding or after its release. Either way, the completeness of all 23S modifications is reassured before mature 70S ribosomes enter the translation pool (70S^RbgA^ and 70S^WT^, Figure 4A).

To confirm the genes involved in the process of h69 modification, five *B. subtilis* protein homologs of the pseudouridine synthase RluD, and methyltransferases RlmH and TlyA were identified using Blast protein search (SI Figure 2). Single-gene deletion mutants of the *B. subtilis* 168 were obtained and characterized relative to the wild-type strain using shotgun LC-MS/MS. Analysis of the rRNA isolates demonstrates that Δ*ylyB* ribosomes lack all three pseudouridine modifications (1940-Ψ,1944-Ψ, 1946-Ψ), Δ*yydA* mutant has no 1944-m^3^Ψ, while both 23S-1949-Cm and 16S-1417-Cm are absent in Δ*yqxC* (SI Figure 7). These observed phenotypes confirm that *ylyB* (NCBI gene ID: 939987), *yydA* (ID: 937752), and *yqxC* (ID: 938651) express homologs of RluD, RlmH, and TlyA (later in text denoted as RluD^h^, RlmH^h^, and TlyA^h^).

To summarize, we assembled a model that incorporates late modification enzymes to the LSU assembly pathway in *B. subtilis (*Figure 4D). Based on the prior findings (52), RbgA acts at the late stage of LSU assembly, where it likely binds to the precursors with a well-developed CP region. Upon binding, RbgA stabilizes several rRNA helices into their mature state (h91-93), thus initiating formation of A and P functional sites. Since RbgA limits access to h69, TlyA^h^ and RluD^h^ must catalyze modification reaction before RbgA binding or after GTP hydrolysis triggers RbgA release. While 70-80% of TlyA^h^ and RluD^h^ dependent modifications are completed by the time 45S-RbgA complex is formed, these modifications may be easily rerouted to take place following RbgA dissociation. Lastly, several r-proteins and other protein factors contribute to LSU maturation including m^3^Ψ methylation by RlmH^h^, which is the final modification step on the assembly pathway.

## DISCUSSION

Using a diverse collection of evidence provided by MS data, a total of 25 modification sites in *B. subtilis* rRNA were identified, and the nucleoside positions and chemistries found agree well with homologous modification sites in other bacteria species (SI Table 1). Out of 25, ten modifications were previously predicted by MS^1^ and RT primer extension analyses (Table 1) and were confirmed here independently using MS^2^ data in coordination with bioinformatics analysis.

A variable modification search implemented in Pytheas can be programmed for placement of virtually any modification types, by specifying their atomic composition, and is fully compatible with custom chemical derivatization and stable isotope labeling. The complexity demonstrated by the inclusion of variable modifications is huge and would be hard to handle manually. Therefore, future developments leading to fast, efficient, and user-friendly software tools for discovery of RNA modifications will benefit the whole MS community to abandon an obviously subjective process of manual MS/MS spectra verification. Access to MS^2^ spectra of modified oligonucleotides from *B. subtilis* ribosome is granted through the PRIDE repository. Given the limited availability of modified oligonucleotide standards, these data might be useful for a better understanding of fragmentation patterns induced by different modification types, for tailoring scoring functions, and further algorithmic developments. Along this line, the Pytheas search used by us is clearly faulty at scoring m^7^G and CMC-Ψ spectra (SI Table 2 and SI Figures 3 and 4C). Due to intrinsic low scores, sequence IDs with m^7^G and CMC-Ψ can easily fall below the overall FDR threshold and modifications can be missed. Work is underway to reflect the needed changes. Another significant drawback of the approach is that only common modification types, previously identified in prokaryotic rRNA, were considered, and if any other non-canonical nucleosides are present, they are left behind by the analyses.

Annotation of the rRNA modification across species is important for understanding their role in translation and antibiotic resistance, but predicting modification sites solely based on the presence or absence of the protein homolog is unreliable. We performed a BLAST protein search across *B. subtilis* proteome (SI Figure 2) to demonstrate that conservation of the modification enzyme may not guarantee conservation of the target site. For example, homologs of *E. coli* methyltransferases RsmC, RsmF, RlmA, RlmG, and RlmI are available, but these enzymes have distinct RNA specificity in *B. subtilis*. Predicting pseudouridine target sites is even more difficult, as pseudouridine synthases from the same enzyme family display a high degree of sequence similarity. To summarize, reliable and conclusive annotation of modifications requires direct experimental evidence.

Although the body of rRNA data collected across bacteria species is limited, the inventory of available modification sites and enzymes ((47) and SI Table 1) suggests that likely there is a core of highly conserved modifications. In *B. subtilis* they correspond to 527-m^7^G, 966-m^2^G, 967-m^5^C, 1402-m^4^Cm, 1498-m^3^U, 1518|1519-m^6^_2_A in 16S, and to 747-m^5^U, 1911-Ψ, 1915-m^3^Ψ, 1917-Ψ, 1939-m^5^U, 2251-Gm, 2445-m^2^G, 2503-m^2^A in 23S (*E. coli* numbering here and below). On the opposite side remain *B. subtilis* specific modifications 1051-m^2^G, 1409-Cm in 16S and 576-m^5^U, 1920-Cm, 2553-Gm, 2574-m^7^G, 2492-Ψ in 23S. Although absent in *E. coli*, most of these nucleosides were identified in other species like *M. tuberculosis* (1409-Cm and 1920-Cm, (25)), *B. stearothermophilus* (2553-Gm, (17)), and in *T. maritima* (1051-m^2^G, (11)). Interestingly, *B. subtilis* 1051-m^2^G found in h34 of 16S is directly opposite to 1207-m^2^G in *E. coli*, which is likely a result of C-m^2^G to m^2^G-C inversion. Furthermore, several novel residues were mapped to the functional sites of the 30S and 50S subunits. Guanosine methylations 2553-Gm (h92), 2574-m^7^G (h90), and pseudouridine 2492-Ψ (h89) belong to the PTC. Cytidine methylations 1409-Cm (16S) and 1920-Cm (23S), also found in several *Mycobacteria* and associated with capreomycin and viomycin sensitivity (25), are part of helices 44 and 69 establishing inter-subunit bridge B2a that connects PTC to the decoding center. The close association with ribosome functional centers suggests that each of these modifications can benefit specific aspects of the *B. subtilis* translation cycle. Unfortunately, the dispensable nature of individual rRNA modification in bacteria makes it technically challenging to link modifications to the specific translational defects that are usually mild (53). As an example, modifications within universally conserved and essential h69, highlighted by us as a result of RbgA depletion, can facilitate processes like initiation, termination, and ribosome recycling, where formation and breakage of the h69-h44 inter-subunit interactions take place (54). Further important to say, that our studies were performed at conditions of steady-state growth in a controlled laboratory environment, and clearly the dynamics of the rRNA modification pattern and stoichiometry may change in response to nutrient availability, stress (55), or to the *B. sublitis* sporulation process.

By examining the LSU assembly pathway that depends on the essential biogenesis factor RbgA we separated 23S modifications into two cohorts. Modifications that are stoichiometrically present in RbgA-dependent 45S intermediate are installed at the early-intermediate stages of assembly, and more studies are needed to further subdivide modification event, determine temporal order and functional dependencies along the pathway. Late modification steps, uncovered by us, include modifications catalyzed by RluD, RlmH, and TlyA homologs. Likely, there is no specific hierarchy between RluD^h^ and TlyA^h^ (and to some degree RbgA) as flexibility of the assembly pathway can provide time for the binding and release of these proteins in an arbitrary order. Based on the cryo-EM structures reported previously for RluD and TlyA associated to their substrates (31,34), enzymes access h69 and flip the base out preparing for the catalysis. RluD (PDB ID: 7BL5) and TlyA (7S0S) have a common protein binding site, and 70-80% of these modifications are completed by the time of RbgA association. Methyl transfer to 1944-m^3^Ψ (*B. subtilis* numbering) depends on prior RluD^h^ pseudouridylation, and RlmH^h^ associates with the late 50S precursors following the release of RbgA (Figure 4D).

Previous studies establish RluD, RlmH, and RlmE ( 2552-Um target site) as late modification enzymes participating in the LSU biogenesis of *E. coli* (36). Unlike RluD and RlmH, RlmE does not have a homolog in *B. subtilis* (SI Figure 2B). Instead, *B. subtilis* utilizes a different enzyme RlmP to form a ribose methylated 2582-Gm (2553-G in *E. coli*) directly adjacent to unmodified 2581-C (2552-Cm in *E. coli*) in h92 stem-loop. According to 45S^RbgA^ inventory (Figure 4A) and the *in vitro* activity of RlmP assayed by Roovers et al. (21), 2582-Gm methylation is complete by the late phases of LSU assembly. Curiously, neither in *E. coli* nor in *B. subtilis,* completion of the modification status strictly follows the order of 23S RNA folding into its canonical conformation. For example, *B. subtilis* 1968-m^5^U situated at h69-h71 junction (Figure 4B) is complete long before two helices are docked. Likewise, 2521-Ψ (h89), 2582-Gm (h92), and 2603-m^7^G (h90) are presented prior to the maturation of the A and P binding sites late on the pathway. While PTC and h69-h71 are known to be the last elements to mature, this further reminds about the complexity of the LSU assembly process in general, and particularly about the lack of understanding of structural rearrangements taking place and additional protein participating in the early assembly phases.

In summary, the provided list of rRNA modification sites leads the way to further discoveries. By understanding the specifics of individual organisms like *B. subtilis* and drawing parallels with other species, including better characterized *E. coli* (SI Table 1 and SI Figure 8), we can gradually define the role of individual RNA modifications and functionally relate modification enzymes to the key steps of the ribosome assembly process. Perhaps, a continued effort of experimental validation of rRNA modification enzymes in *B. subtilis* (18,19,21,22) remains an important direction. By further expanding the list of modification positions across different bacteria species and deciphering the dynamic changes in pattern and stoichiometry they encounter upon stress, limitations, and antibiotic treatment, we will better understand the mechanisms used by RNA modifications to fine-tune translation. Meanwhile, the mass-spectrometry framework and tools for the discovery and quantification of RNA modifications described in this work and the work of others are readily available for accelerating this progress.

## DATA AVAILABILITY

Supplementary Information and Supplementary Data are available for download. RNA mass spectrometry data have been deposited to the ProteomeXchange Consortium via the PRIDE (56) partner repository with the dataset identifiers PXD050367 and PXD051518.

## ACKNOWLEDGEMENTS

We deeply thank the laboratory of Gary Suizdak and particularly Aries Aisporna, Winnie Heim, and Linh Truc Hoang for their help with MS instrumentation and maintenance. We appreciate Julia Polay contribution to the manuscript proofreading process.

---

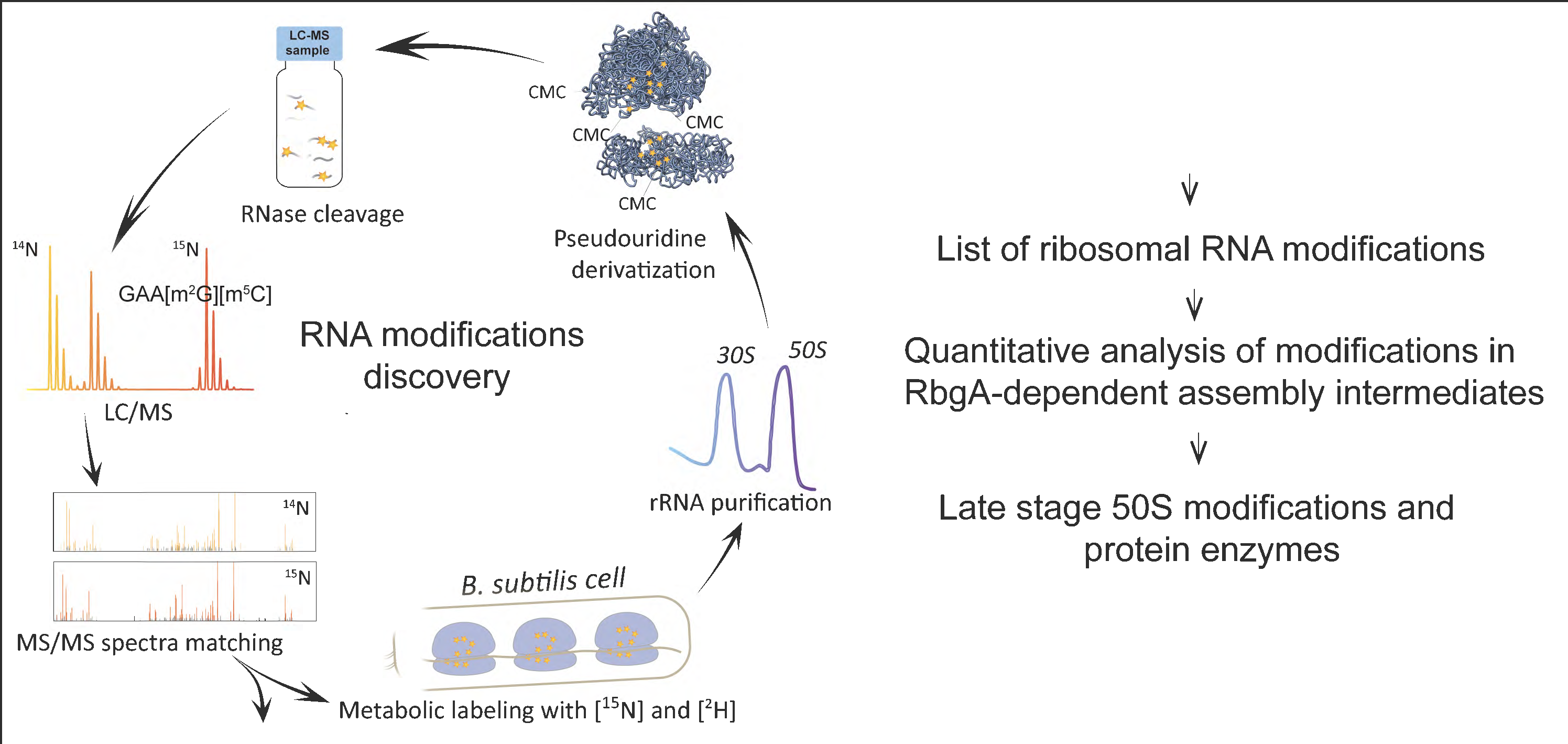

## REFERENCES

1. Sergiev, P.V., Aleksashin, N.A., Chugunova, A.A., Polikanov, Y.S. and Dontsova, O.A. (2018) Structural and evolutionary insights into ribosomal RNA methylation. Nat Chem Biol, 14, 226–235.

2. Taoka, M., Nobe, Y., Yamaki, Y., Sato, K., Ishikawa, H., Izumikawa, K., Yamauchi, Y., Hirota, K., Nakayama, H., Takahashi, N. and Isobe, T. (2018) Landscape of the complete RNA chemical modifications in the human 80S ribosome. Nucleic acids research, 46, 9289–9298.

3. Sergeeva, O.V., Bogdanov, A.A. and Sergiev, P.V. (2015) What do we know about ribosomal RNA methylation in Escherichia coli? Biochimie, 117, 110–118.

4. Sloan, K.E., Warda, A.S., Sharma, S., Entian, K.D., Lafontaine, D.L.J. and Bohnsack, M.T. (2017) Tuning the ribosome: The influence of rRNA modification on eukaryotic ribosome biogenesis and function. RNA Biol, 14, 1138–1152.

5. Watson, Z.L., Ward, F.R., Meheust, R., Ad, O., Schepartz, A., Banfield, J.F. and Cate, J.H. (2020) Structure of the bacterial ribosome at 2 A resolution. Elife, 9.

6. Jeremia, L., Deprez, B.E., Dey, D., Conn, G.L. and Wuest, W.M. (2023) Ribosome-targeting antibiotics and resistance via ribosomal RNA methylation. RSC Med Chem, 14, 624–643.

7. Ofengand, J. (2002) Ribosomal RNA pseudouridines and pseudouridine synthases. FEBS Lett, 514, 17–25.

8. Guymon, R., Pomerantz, S.C., Crain, P.F. and McCloskey, J.A. (2006) Influence of phylogeny on posttranscriptional modification of rRNA in thermophilic prokaryotes: the complete modification map of 16S rRNA of Thermus thermophilus. Biochemistry, 45, 4888–4899.

9. Mengel-Jørgensen, J., Jensen, S.S., Rasmussen, A., Poehlsgaard, J., Iversen, J.J. and Kirpekar, F. (2006) Modifications in Thermus thermophilus 23 S ribosomal RNA are centered in regions of RNA-RNA contact. The Journal of biological chemistry, 281, 22108–22117.

10. Polikanov, Y.S., Melnikov, S.V., Söll, D. and Steitz, T.A. (2015) Structural insights into the role of rRNA modifications in protein synthesis and ribosome assembly. Nat Struct Mol Biol, 22, 342–344.

11. Guymon, R., Pomerantz, S.C., Ison, J.N., Crain, P.F. and McCloskey, J.A. (2007) Post-transcriptional modifications in the small subunit ribosomal RNA from Thermotoga maritima, including presence of a novel modified cytidine. RNA (New York, N.Y, 13, 396–403.

12. Emmerechts, G., Barbé, S., Herdewijn, P., Anné, J. and Rozenski, J. (2007) Post-transcriptional modification mapping in the Clostridium acetobutylicum 16S rRNA by mass spectrometry and reverse transcriptase assays. Nucleic acids research, 35, 3494–3503.

13. Emmerechts, G., Maes, L., Herdewijn, P., Anné, J. and Rozenski, J. (2008) Characterization of the posttranscriptional modifications in Legionella pneumophila small-subunit ribosomal RNA. Chem Biodivers, 5, 2640–2653.

14. Kirpekar, F., Hansen, L.H., Mundus, J., Tryggedsson, S., Teixeira Dos Santos, P., Ntokou, E. and Vester, B. (2018) Mapping of ribosomal 23S ribosomal RNA modifications in Clostridium sporogenes. RNA Biol, 15, 1060–1070.

15. Gutierrez, B., Douthwaite, S. and Gonzalez-Zorn, B. (2013) Indigenous and acquired modifications in the aminoglycoside binding sites of Pseudomonas aeruginosa rRNAs. RNA Biol, 10, 1324–1332.

16. Golubev, A., Fatkhullin, B., Khusainov, I., Jenner, L., Gabdulkhakov, A., Validov, S., Yusupova, G., Yusupov, M. and Usachev, K. (2020) Cryo-EM structure of the ribosome functional complex of the human pathogen Staphylococcus aureus at 3.2 A resolution. FEBS Lett, 594, 3551–3567.

17. Hansen, M.A., Kirpekar, F., Ritterbusch, W. and Vester, B. (2002) Posttranscriptional modifications in the A-loop of 23S rRNAs from selected archaea and eubacteria. RNA (New York, N.Y, 8, 202–213.

18. Nishimura, K., Johansen, S.K., Inaoka, T., Hosaka, T., Tokuyama, S., Tahara, Y., Okamoto, S., Kawamura, F., Douthwaite, S. and Ochi, K. (2007) Identification of the RsmG methyltransferase target as 16S rRNA nucleotide G527 and characterization of Bacillus subtilis rsmG mutants. J Bacteriol, 189, 6068–6073.

19. Desmolaize, B., Fabret, C., Brégeon, D., Rose, S., Grosjean, H. and Douthwaite, S. (2011) A single methyltransferase YefA (RlmCD) catalyses both m5U747 and m5U1939 modifications in Bacillus subtilis 23S rRNA. Nucleic acids research, 39, 9368–9375.

20. Ofengand, J. and Bakin, A. (1997) Mapping to nucleotide resolution of pseudouridine residues in large subunit ribosomal RNAs from representative eukaryotes, prokaryotes, archaebacteria, mitochondria and chloroplasts. Journal of molecular biology, 266, 246–268.

21. Roovers, M., Labar, G., Wolff, P., Feller, A., Van Elder, D., Soin, R., Gueydan, C., Kruys, V. and Droogmans, L. (2022) The Bacillus subtilis open reading frame ysgA encodes the SPOUT methyltransferase RlmP forming 2’-O-methylguanosine at position 2553 in the A-loop of 23S rRNA. RNA (New York, N.Y, 28, 1185–1196.

22. Wolff, P., Labar, G., Lechner, A., Van Elder, D., Soin, R., Gueydan, C., Kruys, V., Droogmans, L. and Roovers, M. (2024) The Bacillus subtilis ywbD gene encodes RlmQ, the 23S rRNA methyltransferase forming m(7)G2574 in the A-site of the peptidyl transferase center. RNA (New York, N.Y, 30, 105–112.

23. Boccaletto, P., Stefaniak, F., Ray, A., Cappannini, A., Mukherjee, S., Purta, E., Kurkowska, M., Shirvanizadeh, N., Destefanis, E., Groza, P. et al. (2022) MODOMICS: a database of RNA modification pathways. 2021 update. Nucleic acids research, 50, D231–D235.

24. Kumar, A., Saigal, K., Malhotra, K., Sinha, K.M. and Taneja, B. (2011) Structural and functional characterization of Rv2966c protein reveals an RsmD-like methyltransferase from Mycobacterium tuberculosis and the role of its N-terminal domain in target recognition. The Journal of biological chemistry, 286, 19652–19661.

25. Johansen, S.K., Maus, C.E., Plikaytis, B.B. and Douthwaite, S. (2006) Capreomycin binds across the ribosomal subunit interface using tlyA-encoded 2’-O-methylations in 16S and 23S rRNAs. Mol Cell, 23, 173–182.

26. Mundus, J., Flyvbjerg, K.F. and Kirpekar, F. (2016) Identification of the methyltransferase targeting C2499 in Deinococcus radiodurans 23S ribosomal RNA. Extremophiles, 20, 91–99.

27. Del Campo, M., Recinos, C., Yanez, G., Pomerantz, S.C., Guymon, R., Crain, P.F., McCloskey, J.A. and Ofengand, J. (2005) Number, position, and significance of the pseudouridines in the large subunit ribosomal RNA of Haloarcula marismortui and Deinococcus radiodurans. RNA (New York, N.Y, 11, 210–219.

28. D’Ascenzo, L., Popova, A.M., Abernathy, S., Sheng, K., Limbach, P.A. and Williamson, J.R. (2022) Pytheas: a software package for the automated analysis of RNA sequences and modifications via tandem mass spectrometry. Nat Commun, 13, 2424.

29. Wolf, E.J., Grunberg, S., Dai, N., Chen, T.H., Roy, B., Yigit, E. and Correa, I.R. (2022) Human RNase 4 improves mRNA sequence characterization by LC-MS/MS. Nucleic acids research, 50, e106.

30. Vanhinsbergh, C.J., Criscuolo, A., Sutton, J.N., Murphy, K., Williamson, A.J.K., Cook, K. and Dickman, M.J. (2022) Characterization and Sequence Mapping of Large RNA and mRNA Therapeutics Using Mass Spectrometry. Anal Chem, 94, 7339–7349.

31. Nikolay, R., Hilal, T., Schmidt, S., Qin, B., Schwefel, D., Vieira-Vieira, C.H., Mielke, T., Burger, J., Loerke, J., Amikura, K. et al. (2021) Snapshots of native pre-50S ribosomes reveal a biogenesis factor network and evolutionary specialization. Mol Cell, 81, 1200–1215 e1209.

32. Stephan, N.C., Ries, A.B., Boehringer, D. and Ban, N. (2021) Structural basis of successive adenosine modifications by the conserved ribosomal methyltransferase KsgA. Nucleic acids research, 49, 6389–6398.

33. Singh, J., Raina, R., Vinothkumar, K.R. and Anand, R. (2022) Decoding the Mechanism of Specific RNA Targeting by Ribosomal Methyltransferases. ACS Chem Biol, 17, 829–839.

34. Laughlin, Z.T., Nandi, S., Dey, D., Zelinskaya, N., Witek, M.A., Srinivas, P., Nguyen, H.A., Kuiper, E.G., Comstock, L.R., Dunham, C.M. and Conn, G.L. (2022) 50S subunit recognition and modification by the Mycobacterium tuberculosis ribosomal RNA methyltransferase TlyA. Proceedings of the National Academy of Sciences of the United States of America, 119, e2120352119.

35. Popova, A.M. and Williamson, J.R. (2014) Quantitative analysis of rRNA modifications using stable isotope labeling and mass spectrometry. Journal of the American Chemical Society, 136, 2058–2069.

36. Rabuck-Gibbons, J.N., Popova, A.M., Greene, E.M., Cervantes, C.F., Lyumkis, D. and Williamson, J.R. (2020) SrmB Rescues Trapped Ribosome Assembly Intermediates. Journal of molecular biology, 432, 978–990.

37. Seffouh, A., Jain, N., Jahagirdar, D., Basu, K., Razi, A., Ni, X., Guarné, A., Britton, R.A. and Ortega, J. (2019) Structural consequences of the interaction of RbgA with a 50S ribosomal subunit assembly intermediate. Nucleic acids research, 47, 10414–10425.

38. Ero, R., Leppik, M., Liiv, A. and Remme, J. (2010) Specificity and kinetics of 23S rRNA modification enzymes RlmH and RluD. RNA (New York, N.Y, 16, 2075–2084.

39. Uicker, W.C., Schaefer, L. and Britton, R.A. (2006) The essential GTPase RbgA (YlqF) is required for 50S ribosome assembly in Bacillus subtilis. Mol Microbiol, 59, 528–540.

40. Jomaa, A., Jain, N., Davis, J.H., Williamson, J.R., Britton, R.A. and Ortega, J. (2014) Functional domains of the 50S subunit mature late in the assembly process. Nucleic acids research, 42, 3419–3435.

41. Sperling, E., Bunner, A.E., Sykes, M.T. and Williamson, J.R. (2008) Quantitative analysis of isotope distributions in proteomic mass spectrometry using least-squares Fourier transform convolution. Anal Chem, 80, 4906–4917.

42. Nakayama, H., Akiyama, M., Taoka, M., Yamauchi, Y., Nobe, Y., Ishikawa, H., Takahashi, N. and Isobe, T. (2009) Ariadne: a database search engine for identification and chemical analysis of RNA using tandem mass spectrometry data. Nucleic acids research, 37, e47.

43. Koo, B.M., Kritikos, G., Farelli, J.D., Todor, H., Tong, K., Kimsey, H., Wapinski, I., Galardini, M., Cabal, A., Peters, J.M. et al. (2017) Construction and Analysis of Two Genome-Scale Deletion Libraries for Bacillus subtilis. Cell Syst, 4, 291–305 e297.

44. Kobayashi, K., Ehrlich, S.D., Albertini, A., Amati, G., Andersen, K.K., Arnaud, M., Asai, K., Ashikaga, S., Aymerich, S., Bessieres, P. et al. (2003) Essential Bacillus subtilis genes. Proceedings of the National Academy of Sciences of the United States of America, 100, 4678–4683.

45. Heiss, M., Reichle, V.F. and Kellner, S. (2017) Observing the fate of tRNA and its modifications by nucleic acid isotope labeling mass spectrometry: NAIL-MS. RNA Biol, 14, 1260–1268.

46. Yan, F., LaMarre, J.M., Rohrich, R., Wiesner, J., Jomaa, H., Mankin, A.S. and Fujimori, D.G. (2010) RlmN and Cfr are radical SAM enzymes involved in methylation of ribosomal RNA. Journal of the American Chemical Society, 132, 3953–3964.

47. Mosquera-Rendon, J., Cardenas-Brito, S., Pineda, J.D., Corredor, M. and Benitez-Paez, A. (2014) Evolutionary and sequence-based relationships in bacterial AdoMet-dependent non-coding RNA methyltransferases. BMC Res Notes, 7, 440.

48. Durairaj, A. and Limbach, P.A. (2008) Improving CMC-derivatization of pseudouridine in RNA for mass spectrometric detection. Anal Chim Acta, 612, 173–181.

49. Addepalli, B. and Limbach, P.A. (2016) Pseudouridine in the Anticodon of Escherichia coli tRNATyr(QPsiA) Is Catalyzed by the Dual Specificity Enzyme RluF. The Journal of biological chemistry, 291, 22327–22337.

50. Zhao, X. and Yu, Y.T. (2004) Detection and quantitation of RNA base modifications. RNA (New York, N.Y, 10, 996–1002.

51. Purta, E., Kaminska, K.H., Kasprzak, J.M., Bujnicki, J.M. and Douthwaite, S. (2008) YbeA is the m3Psi methyltransferase RlmH that targets nucleotide 1915 in 23S rRNA. RNA (New York, N.Y, 14, 2234–2244.

52. Seffouh, A., Trahan, C., Wasi, T., Jain, N., Basu, K., Britton, R.A., Oeffinger, M. and Ortega, J. (2022) RbgA ensures the correct timing in the maturation of the 50S subunits functional sites. Nucleic acids research, 50, 10801–10816.

53. Wang, W., Li, W., Ge, X., Yan, K., Mandava, C.S., Sanyal, S. and Gao, N. (2020) Loss of a single methylation in 23S rRNA delays 50S assembly at multiple late stages and impairs translation initiation and elongation. Proceedings of the National Academy of Sciences of the United States of America, 117, 15609–15619.

54. Ali, I.K., Lancaster, L., Feinberg, J., Joseph, S. and Noller, H.F. (2006) Deletion of a conserved, central ribosomal intersubunit RNA bridge. Mol Cell, 23, 865–874.

55. Fasnacht, M., Gallo, S., Sharma, P., Himmelstoss, M., Limbach, P.A., Willi, J. and Polacek, N. (2022) Dynamic 23S rRNA modification ho5C2501 benefits Escherichia coli under oxidative stress. Nucleic acids research, 50, 473–489.

56. Perez-Riverol, Y., Bai, J., Bandla, C., Garcia-Seisdedos, D., Hewapathirana, S., Kamatchinathan, S., Kundu, D.J., Prakash, A., Frericks-Zipper, A., Eisenacher, M. et al. (2022) The PRIDE database resources in 2022: a hub for mass spectrometry-based proteomics evidences. Nucleic acids research, 50, D543–D552.

